# Spatiotemporal structure of multimodal cues influences color and odor integration in a model insect, the bumblebee *Bombus impatiens*

**DOI:** 10.1101/2024.01.30.577961

**Authors:** Katelyn Graver, Jessica Sommer, Vijay Rao, Giovanni Tafuri, Jordanna D. H. Sprayberry

## Abstract

Bumblebees rely on diverse sensory information to locate flowers while foraging. The majority of research exploring the relationship between visual and olfactory floral cues is performed at local spatial scales and applicable to understanding floral selection. Floral-cue use during search remains underexplored. This study investigates how the bumblebee *Bombus impatiens* uses visual versus olfactory information from flowers across behavioral states and spatial scales. At local spatial scales, non-flying animals in an associative learning paradigm will generalize to either unimodal attribute of a learned color + odor cue with equal likelihood. However, bumblebees flying in a wind tunnel shift cue-use strategy depending on the spatiotemporal scale of cue encounter. When both color and odor cues mimic local/ within patch spatial scale, bumblebees weigh color information of a learned floral-cue more heavily. When cues mimic an intermediate/ between patch spatial scale, bumblebees weigh color and odor information equally, and show the highest response to fully intact multimodal cues. Thus the spatiotemporal scale of sensory information influences how bumblebees utilize multimodal floral cues.

**Statements and Declarations:** This work was internally funded by Muhlenberg College. The authors have no competing interests

## Introduction

Bumblebees are essential pollinators in multiple ecosystems(Heemert et al., 2015; Hegland and Totland, 2008; Klein et al., 2008; Motten, 1986). Human-induced declines in bumblebee populations are therefore cause for concern. In addition to direct factors driving declines (as reviewed by Goulson et al 2015 (Goulson et al., 2015)), anthropogenic activity can indirectly impact the sensory signals that are utilized by bumblebees while foraging for floral resources. For example, negative effects of human activity on floral-scent signaling and reception include a reduction of distance traveled by floral scent (Farré-Armengol et al., 2015; McFrederick et al., 2008), increased foraging times (Fuentes et al., 2016), decreases in floral scent recognition (David et al., 2022; Girling et al., 2013; Sprayberry et al., 2013; Yousry et al.), and alterations of flower visitation rates (Lusebrink et al., 2023; Ryalls et al., 2022). These negative effects are potentially exacerbated by asymmetric impacts of sublethal pesticide exposure, impairing odor-learning but not color-learning (Muth et al., 2019).

However, bumblebees do not exclusively rely on olfactory cues while foraging. Flowers provide a diversity of sensory information to pollinators (e.g., color, shape, patterning, electrostatic fields) (reviewed by Sommer et al. 2022 (Sommer et al., 2022), Rands et al 2023 (Rands et al., 2023)). These signals are information-rich; improving detection, discrimination, and learning (Kulahci et al., 2008; Kunze and Gumbert, 2001; Leonard and Masek, 2014; Leonard et al., 2011). For example, bumblebees have been shown to recognize paired visual and olfactory patterns more efficiently than either cue on its own (Lawson et al., 2017; Leonard et al., 2011). Multimodal signaling can also be used redundantly by experienced foragers, transferring rewarding cues between sensory modalities (Lawson et al., 2017). Learning multimodal floral signals likely improves foraging success for bumblebees, as access to richer floral information reduces uncertainty (Leonard et al., 2011) and improves detection in variable sensory environments (Jordan et al., 2023; Kaczorowski et al., 2012). Understanding the ‘rules’ of multimodal integration of floral information is critical to determining the potential impacts of odor pollution, as these could be amplified or attenuated depending on how scent is integrated with other floral signals. The existing body of work strongly indicates that bumblebees utilize odor information in conjunction with visual cues such as color, but the rules of integration are not fully understood. For example, work by Lawson et al. shows that olfactory pattern learning can be transferred to visual patterns – implying that neural encoding of floral ‘objects’ flexibly integrates multiple modalities – while work by Leonard et al implied that color information dominates over scent (Lawson et al., 2018; Leonard et al., 2011). The variability in the findings and interpretation of color-odor integration studies begets the question: are sensory integration strategies dynamic and, if yes, what foraging contexts trigger a change in integration strategy? Recent work proposed a framework for identifying and understanding foraging phases in terms of state and state-transitions (search *(S)*, acquisition *(A)*, navigation *(N), S*→*A, N*→*A*, etc.), spatial scale (local *(l)*, intermediate *(i)*, distant *(d)*), and forager-background (naive *(n)*, experienced *(e)*, primed *(p)*), with the understanding that different types of sensory information are relevant to different foraging phases (Sommer et al., 2022). Floral information is variable depending on the distance to a flower/ patch. Given that olfactory information is typically resolvable at greater distances from flowers than their visual displays (Sprayberry, 2018), a bumblebee searching for a novel resource might utilize floral information differently than a bumblebee selecting resources within a patch. The majority of multimodal-integration research has been performed at local spatial scales, typically less than or equal to 3.6 meters (reviewed by Sommer et al 2022 (Sommer et al., 2022)). These studies provide valuable data about patch exploitation (*S*→*A*), but their findings may not directly translate to bumblebees in the search (S) or navigation (N) phases. Given that bumblebees forage over large distances, up to 1 kilometer (Hellwig and Frankl, 2000; Lihoreau et al., 2012; Osborne et al., 1999), an understanding of how spatial scale and behavioral state influence multimodal integration is crucial to furthering our understanding of foraging behavior. This phenomenon is not isolated to bumblebees, but applies to many species of central place foragers (Sommer et al., 2022). The study presented here uses a combination of associative odor learning and wind tunnel paradigms to ask if sensory integration strategies are flexible depending upon the (simulated) foraging phase (Tables 1, 2). Specifically, we use the free-moving proboscis-extension-reflex paradigm (Muth et al., 2017; Sprayberry, 2020) to ask if bumblebees trained to a multimodal color+odor cue will generalize to the learned color paired with a novel odor and vice versa (State(scale, background) = *A(l,e)*). We use a wind-tunnel paradigm to ask how foraging bumblebees trained to a multimodal color+odor ‘flower’ will respond to flowers with varying trained/ novel color + odor values in a landing assay. In addition, we manipulate access to visual and olfactory cues to simulate changes in spatial scale: tests where bees only have access to odor cues at the launch point are intended to mimic an intermediate distance from a flower (*N*→*A(i,e)*), while those where both color and odor cues are available represent local spatial scales (*N*→*A(l,e)*) (Sprayberry, 2018).

**Table 1.** Experimental conditions for FMPER experiments. Testing conditions that are congruent with training are highlighted green and marked with a check. Testing conditions that are novel are highlighted grey and marked with an X.

| Question Asked | Associative Condition |  | Test Stimulus Conditions |  |  |  |
| --- | --- | --- | --- | --- | --- | --- |
|  |  |  | Choice 1 |  | Choice 2 |  |
|  | color | odor | color | odor | color | odor |
| Is a multimodal cue learned? | blue | LoV | blue(✓) | LoV(✓) | yellow(X) | JB(X) |
|  | yellow | LoV | yellow(✓) | LoV(✓) | blue(X) | JB(X) |
| Does color or odor exhibit dominance in multimodal learning? | blue | LoV | blue(✓) | JB(X) | yellow(X) | LoV(✓) |
|  | yellow | LoV | yellow(✓) | JB(X) | blue(X) | LoV(✓) |
| Is accurate odor with dissonant color generalized? | blue | LoV | yellow(X) | LoV(✓) | yellow(X) | JB(X) |
|  | yellow | LoV | blue(X) | LoV(✓) | blue(X) | JB(X) |
| Is accurate color with dissonant odor generalized? | blue | LoV | blue(✓) | JB(X) | yellow(X) | JB(X) |
|  | blue | JB | blue(✓) | LoV(X) | yellow(X) | LoV(X) |
|  | yellow | LoV | yellow(✓) | JB(X) | blue(X) | JB(X) |
|  | yellow | JB | yellow(✓) | LoV(X) | blue(X) | LoV(X) |

**Table 2.** >Wind tunnel experimental conditions. All bees were trained to a blue + lily of the valley (LoV) scented flower. Congruent testing conditions are green highlighted and marked with a check. Incongruent conditions are grey highlighted and marked with an x.

| Wind Tunnel Set-up | Test Flower Condition |  |
| --- | --- | --- |
|  | color | odor |
| opaque panel | blue (✓) | LoV(✓) |
|  | blue(✓) | JB(X) |
|  | yellow(X) | LoV(✓) |
|  | yellow(X) | JB(X) |
| clear panel | blue(✓) | LoV(✓) |
|  | blue(✓) | JB(X) |
|  | yellow(X) | LoV(✓) |
|  | yellow(X) | JB(X) |

## Methods

### Animals

This study used 13 colonies of *Bombus impatiens* (Koppert Biological Systems or BioBest) between 2018 and 2023. The temperature of the colonies was maintained at 24–29° C by wrapping a seedling heat map with an attached thermostat around each. All colonies had continued access to ad-libitum pollen throughout the studies. Colonies used for free-moving proboscis-extension-reflex (FMPER) experiments had ad-libitum access to 30% sucrose solution, while those used in wind-tunnel experiments only had access to sucrose during a daily feeding window of 2-3 hours to increase experimenter access to foraging bees for experimental-trials.

#### Stimulus Modalities

All experiments tested color and scent sensory modalities. The colors used were yellow and blue (Ings et al., 2009); scents were lily of the valley (LoV) and juniper berry (JB) (Sprayberry, 2020). FMPER experiments were performed with strips cut from plastic office folders (Office Depot), and wind-tunnel experiments used 3-D printed flowers (eSun PLA filament). Conditioning for FMPER experiments is rapid enough that multiple associative color+scent conditions could be tested, while all wind tunnel experiments were based on association to blue+LoV-scented flowers (diameter 8.9 cm). Responses to combinations of novel and congruent combinations of trained modalities were then tested with different experimental paradigms to determine if behavioral state impacts the generalization of learned multimodal information (FMPER, Table 1; wind tunnel, Table 2).

#### Localized Associative Reward Learning: FMPER

Associative learning was assessed using Free Moving Proboscis Extension Reflex (FMPER) (Muth et al., 2017; Sprayberry, 2020). Active individual *B. impatiens* were selected from lab colonies and placed into screen-backed vials to acclimate for 2 hours. During experiments the vials were placed into a ventilating FMPER apparatus (Fig 1). The ventilation system drew air in through two small holes in the vial lid and out of the back screens, with flow rates ranging between 0.1 to 0.3 m/s (VWR-21800-024 hot wire anemometer). During testing bees were able to walk back and forth in the vials, but could not fly and had a relatively small range of motion.

**Fig 1.**
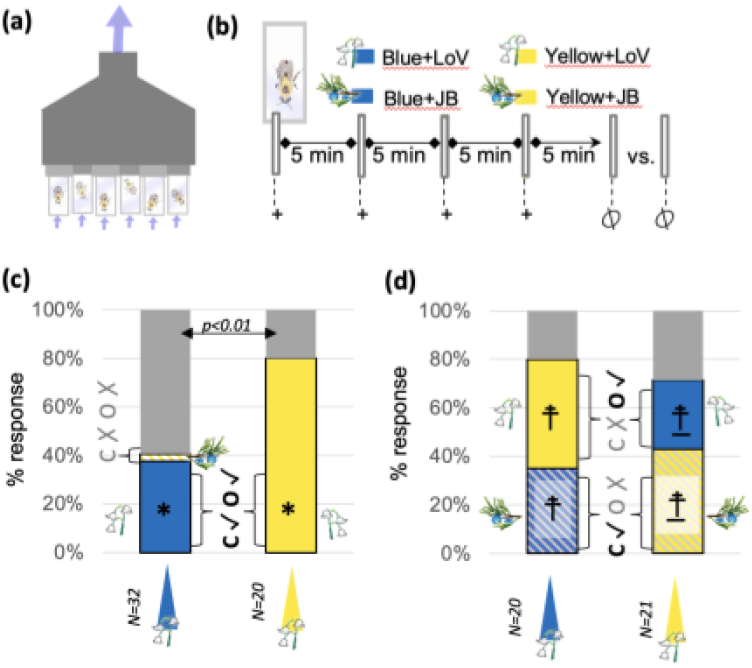
Free-moving proboscis extension reflex (PER) experiments use absolute conditioning to explore multimodal (color+odor) cue learning. A) A ventilating array holds six bees in screen backed tubes, allowing for multiple bees to be tested. B) Tube-lids have two holes, each of which allow a scented+ colored strip to be inserted. The odor was lily of the valley (LoV) or juniper berry (JB) and the color was blue or yellow; the rectangles in the legend show the color condition as their fill color, and the scent condition as their outline color (LoV = pink and JB = green). This color schema is consistent throughout all figures. For the four training trials the strips contain a drop of 50% sucrose. For test trials two unrewarding strips were inserted. If bumblebees extended their proboscis onto a strip that choice was recorded and both strips were removed. If bees did not make a choice, a “no choice” response was recorded. All trials (training and testing) were separated by approximately 5 minutes. C&D) training conditions are indicated by colonng of triangles below response distributions, while colors within the response distribution indicate choice frequencies for test conditions. No choice responses are plotted in grey, with arrows indicated p values of no-choice comparisons across treatments.. Congruous testing cue components are market with a ✓, while dissonant components are marked with an ✗. Symbols on response-choices denote p values for binomial comparisons of choices: *p<0.001, †p>0.2. An underscore (_) indicates a confidence interval > 0.5, due to a small number of bees making a choice. C) Bumblebees will choose a learned multimodal cue more frequently than a completely dissonant cue. D) Both odor and color are generalized by bumblebees, with neither cue showing dominance.

During conditioning, individual bees were offered a single drop of 50% sucrose on either a blue or yellow plastic strip that was inserted through one of the holes in the vial lid (hole selection was randomized). Absorbent adhesive bandage tape (Cover Roll) was placed on the back of the strips in order to hold an odor-stimulus - 1 uL of either Lily of the Valley (LoV) or Juniper Berry (JB), depending on the condition being tested. Bumblebees would undergo four association trials, during which they were presented with these rewarding color+scent strips at 5-minute intertrial intervals. Individuals that actively participated in the four association trials would then undergo a test trial after an additional 5-minute wait. During tests bees were presented with two unrewarding color+scent strips; the sensory values were variable based on trial condition. Trials were designed to answer four different questions: 1) is a multimodal cue learned in this paradigm?; 2) does color or odor exhibit dominance in this paradigm?; 3) is accurate odor with dissonant color generalized?; 4) is accurate color with dissonant odor generalized? (Table 1). Proboscis extension onto a strip was considered a choice and individuals that approached the strips three times or went 45 seconds without exhibiting PER were considered no-choice. All tested bees were tagged following experiments to prevent re-testing and ensure statistical independence of data points.

#### Wind Tunnel Tests

##### Association

During daily timed feeding sessions, wind-tunnel testing colonies were given access to a glass feeder with a LoV-scented blue flower/s (one or two) containing 30% sucrose. Both the feeder base and lowers were 3-D printed with non-toxic PLA filament (eSun blue). Flower shape was based on the silhouette of an *Echinacea purpurea* flower, with a rectangular hole in the center that fit onto the nectar outlet of the feeder base. The print file is provided in the supplemental materials. Odor stimuli were added by pipetting 5 uL of the LoV essential oil onto a 2 cm x 2 cm piece of absorbent tape (Cover Bandage) on the back of each flower. After three consecutive days of flower color + scent exposure, colonies were eligible for wind tunnel trials. Occasionally throughout a colony’s lifespan, the foraging activity within the timed feeding window was not robust enough to maintain adequate levels of honey for the hive. In these cases ad libitum feeding was offered without floral stimuli for two to three days. Following this, colonies would have an additional two days to re-associate with the color and scent stimuli. Following timed feeding sessions, 3D-printed flowers were washed and aired out overnight.

##### Wind tunnel structure

The wind tunnel dimensions were 1.7 m x 0.5 m x 0.47 m; the walls and floor were white corrugated plastic and the ceiling was transparent plexi-glass. Green tape ran lengthwise on the walls, floor, and ceiling to provide visual cues of the wall locations without providing widefield field motion cues for distance measurement by flying bees (Baird et al., 2021). Air flow through the tunnel is provided by a box fan, with insulation and white plastic egg-crate light-diffuser ceiling panels (Plaskolite) to even out flow. Panels were placed at ⅓ and ⅔ of the way through the tunnel on opposite sides, making a serpentine maze (Figure 2). Air flow through the tunnel is not laminar, but glycerine smoke-plumes were visualized to confirm that scent plumes are carried all the way through the tunnel. Average air flow sampled in the center of the tunnel in each section was 0.37 +-0.11 m/s. Test bees were introduced into the tunnel at a consistent location, via a launch platform that accepts bee-vials. Test flowers were placed at the opposite end of the tunnel, 1.5m from the launch platform.

#### Testing

Bees that were seen actively feeding from the training flowers in the colony were removed and placed in plastic vials for wind tunnel experimentation, waiting on a seedling heating mat (Propagate Pro, 27-28 C) for 15-40 minutes before being placed on the launch platform of the wind tunnel. The cap to the bee’s vial was removed so that the bee could exit the vial, and this cap removal officially began a trial. Bees that did not exit their vial within five minutes were excluded from analysis. Bees that exited the vial were given 10 minutes to locate and/or land on the test flower. Test flowers were unrewarded 3D-printed copies of the training flowers (either blue or yellow), scented with 10 uL of odor (1:100 LoV or 1:100 JB; diluted in unscented carrier), depending on the experimental condition being tested (Table 2). Bees that landed on the test flower were given a “land” designation and the time in which they landed on the flower was recorded. Bees that did not choose to land after 10 minutes were removed from the tunnel and marked as “no-land.” In addition, we recorded if bees navigated upwind to the third section of the wind tunnel. All tested bees were tagged and returned to their colony following experimentation to prevent re-selection and testing, ensuring statistical independence of data points.

#### Testing effect of cue encounter

Previous work indicates that searching bumblebees most likely encounter floral cues as either odor first, or odor and visual cues simultaneously (Sprayberry, 2018). In an effort to push bumblebees into a searching behavioral state (Sommer 2022), we placed panels at ⅓ and ⅔ of the way through the wind tunnel on opposite sides to create a serpentine-maze path. These tests endeavor to investigate search behavior in bumblebees at both a local spatial scale, with clear panels allowing access to both visual and odor cues from the launch point, and an intermediate spatial scale, with opaque panels obscuring visual cues (Figure 3). We used the number of tested individuals that traveled to the third section to calculate the percentage of upwind-flight bees. We then used the number of individuals who landed divided by the number of upwind-flying bees to calculate the percent landing.

**Fig 3.**
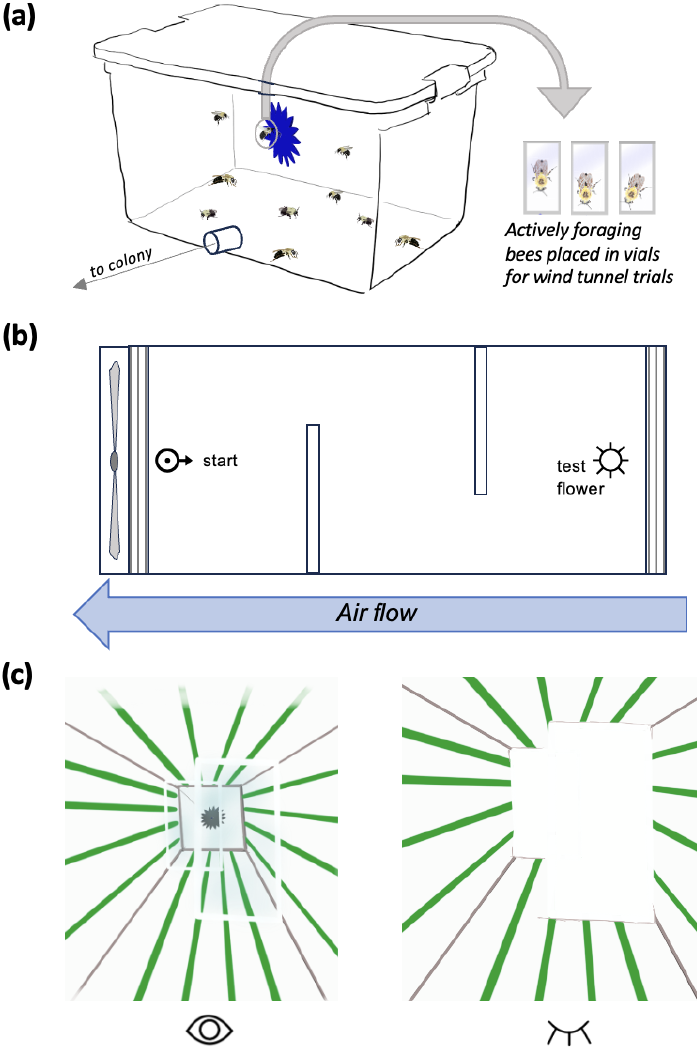
(a) Wind tunnel experiments select actively foraging bumblebees from the colonly’s training flower and set them in vials for wind tunnel trials, (b) The wind tunnel has two panels that break it into a serpentine path over three sections between the start and an unrewarding test flower. Bumblebee vials are placed at the ‘start’ of the wind tunnel. After vials are uncapped bees have 10 minutes to complete the wind tunnel maze. Data on which bees navigate to the third section, and which land on the test flower are recorded, (c) The panels that divide the wind tunnel are either clear, allowing a starting bee to both see and smell the test flower, or opaque, resulting in the starting bee only having access to scent cues.

#### Statistical Analysis

##### FMPER Analysis

For each experimental question analysis of trial data proceeded as follows: 1) Full response distributions (including no choice) were tested against random (33%) with a fisher’s exact test. 2) Correct/ incorrect choices were tested against random (50%) with a binomial distribution test. 3) No choice data were tested with pairwise Fisher comparisons (using the pairwise.fisher.test function in R). Alpha values for the first two tests were Holm-Bonferroni corrected. P-values for the third test were corrected with the Benjamani-Hockberg method. All analyses are summarized in Table 3.

**Table 3.** >Statistics for FMPER experiments.

| Question Asked | Associative Condition | test against random distribution |  |  | test for correct responses > incorrect responses |  |  |  | test for impact of associative condition in no-choice responses |  |
| --- | --- | --- | --- | --- | --- | --- | --- | --- | --- | --- |
|  |  | p value | Holm Thresh. | n | p value | Holm Thresh. | delta CI | n | comparison | p value |
| Is a multimodal cue learned? | blue-LoV | 0.00006 | 0.025 | 32 | 0.002 | 0.05 | 0.32 | 13 | B-LoV vs Y-LoV | 0.009 |
|  | yellow-LoV | 0.00001 | 0.025 | 20 | 0.00003 | 0.05 | 0.21 | 16 |  |  |
| Does color or odor exhibit dominance in multimodal | blue-LoV | 0.43 | 0.025 | 20 | 0.44 | 0.05 | 0.5 | 16 | B-LoV vs Y-LoV | 0.72 |
|  | yellow-LoV | 0.77 | 0.05 | 21 | 0.6 | 0.025 | 0.52 | 15 |  |  |
| Is accurate odor with dissonant color generalized? | blue-LoV | 0.02 | 0.025 | 19 | 0.02 | 0.05 | 0.44 | 10 | B-LoV vs Y-LoV | 0.54 |
|  | yellow-LoV | 0.01 | 0.05 | 21 | 0.09 | 0.025 | 0.55 | 9 |  |  |
| Is accurate color with dissonant odor generalized? | blue-LoV | 0.002 | 0.025 | 20 | 0.11 | 0.05 | 0.58 | 6 | B-LoV vs. B-JB | 0.28 |
|  |  |  |  |  |  |  |  |  | B-LoV vs Y-LoV | 0.002 |
|  |  |  |  |  |  |  |  |  | B-LoV vs Y-JB | 0.75 |
|  | blue-JB | 0.0001 | n/a | 26 | 1 | 0.025 | 0.24 | 13 | B-JB vs Y-LoV | 0.004 |
|  |  |  |  |  |  |  |  |  | B-JB vs Y-JB | <0.001 |
|  | yellow-LoV | 0.00001 | 0.025 | 23 | 0.0001 | 0.05 | 0.17 | 19 | Y-LoV vs Y-JB | 0.000008 |
|  | yellow-JB | 0.00000006 | 0.025 | 32 | 0.004 | 0.05 | 0.31 | 8 |  |  |

##### Wind Tunnel Analysis

Both upwind-flight and land data were analyzed as no-choice data above. For each, data for all stimulus conditions within a set of wind tunnel trials (opaque or clear maze-dividers) were tested with pairwise Fisher comparisons (using the pairwise.fisher.test function in R). P-values were corrected with the Benjamani-Hockberg method. All analyses are summarized in Table 4. Land data were further analyzed with two different logistic regression models (using the glm function in R): one ‘component’ model with separate variables for color and odor to represent assessing sensory variables independently, and one ‘synthetic’ model with a single ‘color + odor match’ variable to represent assessing sensory variables as an amalgam. Both models were run on both panel conditions (clear and opaque). The AIC was calculated for each model run and used to determine the relative likelihood between models.

**Table 4.** Wind Tunnel experiment statistics.

| Panel Condition | Test Flower | Upwind Flight: comparison p-values |  |  |  | Landing: comparison p-values |  |  |  |
| --- | --- | --- | --- | --- | --- | --- | --- | --- | --- |
|  |  | n | yellow-LoV | blue-JB | yellow-JB | n | yellow-LoV | blue-JB | yellow-JB |
| clear | blue-LoV | 20 | 1 | 0.34 | 1 | 12 | **0.02** | 0.24 | **<0.001** |
|  | blue-JB | 18 | 0.34 | - | 0.34 | 15 | 0.17 | - | **0.002** |
|  | yellow-LoV | 25 | - | 0.34 | 1 | 14 | - | 0.17 | 0.15 |
|  | yellow-JB | 21 | 1 | 0.34 | - | 13 | 0.15 | **0.002** | - |
| opaque | blue-LoV | 18 | 0.92 | 0.92 | 1 | 12 | **0.001** | **0.003** | **<0.001** |
|  | blue-JB | 16 | 0.92 | - | 0.92 | 12 | 1 | - | 0.5 |
|  | yellow-LoV | 25 | - | 0.92 | 0.92 | 15 | - | 1 | 0.5 |
|  | yellow-JB | 20 | 0.92 | 0.92 | - | 13 | 0.5 | 0.5 | - |

## Results

### Non-foraging bumblebees will utilize individual components of a learned multimodal cue

To test bumblebees’ ability to learn a color+odor cue, we trained bees to associate a scented-color strip with a sucrose reward using the free-moving proboscis extension reflex paradigm (FMPER), which tests for proboscis extension reflex (PER) after conditioning using two unrewarding strips: one matching their training condition, and one with dissonant color+odor (Table 1, Fig 1). In both training conditions (blue+lily of the valley (LoV) and yellow+LoV) the trained cue was selected more frequently than the dissonant cue (p<0.01, Table 3). We then tested if either the color or odor component of a learned color+odor cue was dominant by testing trained bees with a choice of learned-color + novel-odor versus novel-color + learned-odor stimuli. In these tests bumblebees were equally likely to select the learned-color or learned-odor (Fig 1d, Table 3). We also tested if bumblebees would generalize to individual learned components of a color+odor cue when compared against an entirely novel cue. This was performed for each sensory modality, where bumblebees trained to a color+odor cue were tested against the learned-color+novel-odor in one set of experiments, and against the learned-odor+novel-color in another set (Figure 2). Bumblebees did generalize to a learned cue component for at least one associative condition in each modality: bees trained to blue+LoV selected yellow+LoV at a higher frequency than yellow+juniper berry (JB); and bees trained to yellow-LoV selected yellow+JB at a higher frequency than blue+JB (Figure 2, Table 3).

However, the responsiveness of bumblebees within this FMPER paradigm was variable based on the color and odor values of multimodal cues (Table 3). Bumblebees associated to blue-strips demonstrated a higher percentage of no-choice responses than those associated to yellow-strips in tests of multimodal learning ability (p=0.009, Table 3). Experiments on bumblebee likelihood of generalizing to a learned color + novel odor show similar trends; all tests utilizing blue associative strips have a higher percentage of no-choice responses (Table 3). These tests also show an increase in no-choice when the associative odor is juniper berry. Indeed, color generalization trials where bumblebees were associated to blue+JB not only showed a high percentage of no-choice responses, but bees did not generalize to the learned color. Rather they chose yellow+LoV in their test phase. Likewise, bees trained on yellow+LoV had the highest percentage of generalization responses; and bees trained to yellow+JB showed a significant increase in no-choice responses (Figure 2).

### Temporal structure of cue encounter correlates with a shift in how foraging bumblebees use multimodal information

To test how actively foraging bumblebees respond to components of learned multimodal cues, we first trained colonies to forage on blue-LoV scented flower feeders. Foraging bees were then collected from feeders and flown in a wind tunnel divided into a three-section maze with offset panels; the final section held an unrewarding flower. We used the number of tested individuals that traveled to the third section to calculate the percentage of upwind-flying bees. To filter out non-responsive bees, we then used the number of upwind-flying bees who landed on the test flower to calculate the percent landing (Figure 3). Wind tunnel experiments measured the responses of actively foraging bees to various novel or congruent combinations of color and odor (Table 2). The percentage of upwind-flying bees was consistent across all treatments; unaffected by panel-status, color, or scent (p>0.3, Figure 4, Table 4). However, cue-value had a substantial effect on landing frequency, with the blue-LoV flowers having the highest landing frequency, and flowers with any dissonant modalities showing reduced frequency in most conditions (Figure 4, Table 4).

**Figure 4.**
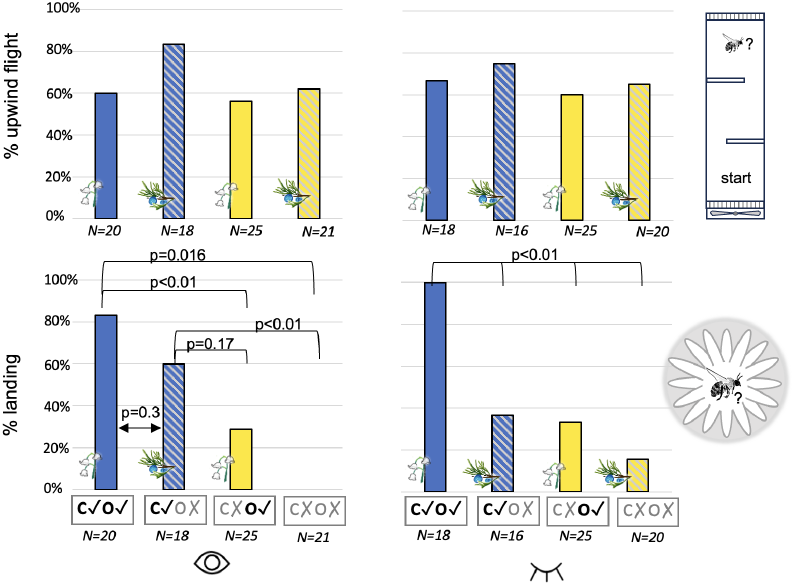
Foraging bumblebees flown in the wind tunnel utilize multimodal cue information differently depending on availability of cues at takeoff. All bees in these experiments were associated to a blue-LoV scented flower. The top row of graphs show the percentage of tested bees that flew upwind to the third wind tunnel section (% upwind flight). The bottom row of graphs shows the percentage of navigating bees that landed on an unrewarding test flower (% landing). Test flower conditions varied; congruous testing cue components are marked with a ✓, while dissonant components are marked with an ✗. The left-hand column of plots show data for wind tunnel tests with clear panels, so that both visual and olfactory information was available at launch. The right-hand column of plots show data for tests with opaque panels, so that only odor information is available at launch. All p-values shown are from pairwise Fisher comparison tests and corrected with the Benjamani-Hockberg method.

Wind tunnel tests endeavor to investigate behavior in bumblebees across spatial scales: clear panels facilitate simultaneous visual and odor cue encounter to simulate a local-spatial scale; while opaque panels create a temporal offset, with odor preceding visual cue encounter, to simulate an intermediate spatial scale (Figure 3) (Sommer et al., 2022; Sprayberry, 2018). When bees encountered odor cues before visual cues they showed bimodal landing responses; responding robustly to only the learned multimodal cue combination blue+LoV (Figure 4). Bees that encountered visual cues synchronously with odor showed a gradient of landing frequencies; with the highest response to learned-color + learned-odor cues, followed by learned-color + novel-odor, then novel-color + learned-odor, and no response to a fully novel cue (Figure 4). Logistic regression analysis confirmed the differential use of color and odor information across the two panel conditions. While a component model (where variables are flower color and flower odor) was equally predictive of responses in both conditions, the synthetic model (with a single variable representing combined flower color and odor information) had divergent responses across conditions: clear panel data were poorly represented with the synthetic model, while opaque panel data were better predicted (Table 5). A comparison of component model results for the two panel conditions offers some insight into this difference. For clear panel/ simultaneous cue-encounter trials, the coefficient for color is almost double that of the odor coefficient, resulting in a higher probability of landing for congruent color + novel odor cues compared to novel color + congruent odor (Table 6, Fig 5). For opaque panel/ odor-cue primed trials, the color and odor coefficients are equal, resulting in equal landing probability for combined congruent+novel cues (Table 6, Fig 5).

**Table 5.** Statistics for two different logistical models. The ‘component’ model separates cue conditions into visual (match, no match) and olfactory (match, no match). The ‘synthetic’.

| Panel Condition | Component Model |  |  | Synthetic Model |  |  | Likelihood Ratio |
| --- | --- | --- | --- | --- | --- | --- | --- |
|  | R2 | p | AIC | R2 | p | AIC |  |
| Clear | 0.33 | 5.0E-06 | 56 | 0.15 | 0.001 | 67 | 0.004 |
| Opaque | 0.3 | 3.0E-05 | 56 | 0.34 | 1.0E-06 | 51 | 0.08 |

**Table 6.** Coefficients for the component model in both experimental conditions. The spatiotemporal structure of cue encounter influences the relative weighting of color and odor information.

|  | Odor Primed/ Opaque Panel |  | Simultaneous Cue/ Clear Panel |  |
| --- | --- | --- | --- | --- |
|  | coefficient | p-value | coefficient | p-value |
| intercept | -2.6 | 0.001 | -3 | 0.0007 |
| color | 2.6 | 0.002 | 3.2 | 0.0002 |
| odor | 2.3 | 0.005 | 1.8 | 0.03 |

## Discussion

### The spatiotemporal scale of cue encounter modulates usage of multimodal information

Previous multimodal studies on bumblebees have shown support for the ‘efficacy backup hypothesis’, where access to information across multiple sensory modalities improves learning and recall in noisy environments (Kaczorowski et al., 2012; Lawson et al., 2017; Lawson et al., 2018; Leonard et al., 2011). The data presented here also support this hypothesis; with our findings that unimodal components from a learned multimodal cue are capable of influencing behavioral choices in our associative learning FMPER paradigm (figs 1,2). In addition, a substantial percentage of bees flown in the clear paneled wind tunnel were responsive to the learned color component of trained floral cues when paired with a dissonant odor value (fig 4), a finding that matches foundational work on color-odor integration in bumblebees (Leonard et al., 2011). From a neural circuitry perspective, this implies bumblebees are able to access and evaluate the unimodal components of learned multimodal cues during the activation of behavioral responses. However, bumblebee responses to unimodal cue-components paired with dissonant values are not uniform across experimental paradigms. Bumblebees flown in an opaque-paneled wind tunnel, intended to simulate an experienced forager navigating at an intermediate spatial scale (foraging state *N*→*A(i,e)*), showed the highest landing frequencies on flowers with fully intact multimodal cues. The principal difference between wind tunnel paradigms is the temporal pattern of sensory encounters. Opaque panels allow access to odor cues *before* color; indeed, this phased cue encounter is how ‘intermediate’ spatial scales in central place foragers have been defined (Sommer et al., 2022). Clear panels facilitate either simultaneous color+odor or color cue encounter first, simulating experienced foragers navigating at local spatial scales (foraging state *N*→*A(l,e)*). The neural mechanisms behind how the timing of multimodal cues can modulate the relationship between unimodal cues have received little attention. A recent study utilizing conditioning of the proboscis extension reflex in harnessed bumblebees investigated the impact of color and odor synchronicity on bumblebee learning (Riveros, 2023). In their experimental context, they also found evidence the bumblebees would generalize to an unimodal component of a learned bimodal; with readily learned cue combinations showing stronger responses to odor alone. Interestingly bees trained with odor preceding color showed the lowest responses to unimodal cues, providing some support for hypothesizing that the temporal structure of odor versus color encounter could modulate how bumble bees process multimodal information.

However, that treatment was also poorly learned in their experimental paradigm thus the mechanisms behind spatiotemporal scales of multimodal cues and behavioral activation remain an intriguing site for future study.

This study uses spatiotemporal structure to simulate different free-foraging contexts, which should modulate a forager’s ‘state’. The idea that state influences sensory processing in insects is not novel. Studies on fruit flies, hawk moths, and mosquitoes have explored how shifts in state can underlie behavioral flexibility (Ache et al., 2019; Chapman et al., 2018; van Breugel et al., 2015; Vogt et al., 2021). Flight activity is known to modulate sensory processing - enhancing visual processing of landing in fruit flies (Ache et al., 2019) and olfactory processing in hawk moths (Chapman et al., 2018). Hunger can modulate responses to odor stimuli - converting aversion to attraction - in fruit flies (Vogt et al., 2021). Similarly, in mosquitoes, one sensory cue (CO_2_) can influence responses to an additional cue (host odors) (van Breugel et al., 2015). In the current body of literature, the presence of one sensory cue (internal or external) modulates the responses to an additional cue. The results presented here show modulation based on the spatiotemporal structure of the same suite of external sensory cues; which is a novel, complex instance of state-modulated sensory processing.

### Ecological implications and limitations

Understanding how the multimodal information flowers provide is utilized by bumblebees in different foraging phases and across gradients of sensory disruption is critical to conservation efforts. For example, if a learned color paired with a novel odor is generalized to the learned multimodal cue by a substantial portion of within-patch foragers - as indicated by our clear-panel wind tunnel tests - then local disruption of floral odors via anthropogenic activities may be ameliorated. However, generalization tendencies are likely impacted by the valence and foraging state of any odor disruptions. In other words, if odors are *aversive* rather than just different, we may not see the same generalization tendencies. For example, Ryalls et al found that diesel pollution on a local level substantially reduced the visitation of a broad spectrum of pollinators, including bumblebees (Ryalls et al., 2022). A subsequent study found variable effects of diesel-driven odor pollution dependent on floral color (Lusebrink et al., 2023). Interestingly diesel exhaust contains sulfurous components - an attribute shared by several fungicides that have also been shown to disrupt bumblebee behavior on a local scale (David et al., 2022; Sprayberry et al., 2013). Thus while our current findings offer hope that odor pollution may not be problematic at local spatial scales, a more complete understanding of odor valence is needed before dismissing concerns entirely.

When considering bumblebees searching for resources at a larger spatial scale, the impacts of anthropogenic disruption are potentially more dire. Bumblebees travel large distances from their nest during foraging bouts (Hellwig and Frankl, 2000; Lihoreau et al., 2012; Osborne et al., 1999), presumably searching for resources if they are not navigating directly to a previously learned patch. Recent work showed that bumblebees prefer to fly upwind (Combes et al., 2023), which primes them to utilize olfactory navigation if they encounter an ‘acceptable’ odor plume (Baker et al., 2018; Carde and Willis, 2008). If, as our opaque-panel tests indicate, experienced foraging bees are less flexible in terms of accepting disrupted color+odor combinations they are more likely to pass over discovered patches whose odors have been modulated by human activity. Given that both habitat loss and fragmentation reduce total resources available to bumblebees and increase the distance between viable resource patches (reviewed by Goulson (Goulson et al., 2015)), anthropogenic-odor disruption could result in consequential reductions in foraging efficiency. Indeed, understanding the impact of spatial scale on odor generalization within multimodal cues is a critical next step in determining the conservation impacts of odor pollution.

## Acknowledgements

The authors would like to thank Muhlenberg College for financial support of this work through summer research stipends. We thank N. Defino, K. Esbenshade, N. Yousry, and P. Henderson for contributions to equipment construction and data collection.

## Ethics

Research on insects does not require approval by IACUC (Institutional Animal Care and Use Committee) in the United States, thus no formalized ethical approval is available for this study. However, my lab historically and currently adheres to the “3Rs” laid out in Crump et al’s 2023 “Is it time for insect researchers to consider their subjects’ welfare?”(Crump et al., 2023).

## Data Availability

All data will be available for download as a supplementary file (S1) when published in a peer-reviewed journal.

